# Decoding functional proteome information in model organisms using protein language models

**DOI:** 10.1101/2024.02.14.580341

**Authors:** Israel Barrios-Núñez, Gemma I. Martínez-Redondo, Patricia Medina-Burgos, Ildefonso Cases, Rosa Fernández, Ana M. Rojas

## Abstract

Protein language models have been tested and proved to be reliable when used on curated datasets but have not yet been applied to full proteomes. Accordingly, we tested how two different machine learning based methods performed when decoding functional information from the proteomes of selected model organisms. We found that protein Language Models are more precise and informative than Deep Learning methods for all the species tested and across the three gene ontologies studied, and that they better recover functional information from transcriptomics experiments. The results obtained indicate that these Language Models are likely to be suitable for large scale annotation and downstream analyses, and we recommend a guide for their use.

## INTRODUCTION

The current era of genomic biodiversity is associated with the generation of thousands of sequences, with around 250 million protein sequences now registered in the January release of UniProtKB. However, it is all too common that the function of these sequences is not determined experimentally. The infamous gap between experimental data and the number of sequences available has remained unbridged for several decades now, with experimental annotations associated with less than 0.5% of known proteins (1). At present, functional annotation is most often carried out through computational prediction, relying inherently on evolutionary information. Moreover, studying the functional role of proteins is commonly biased by the particular interests of certain areas of the scientific community, such that the comprehensive study of most genes is rarely attempted. For instance, most research in humans was done on a small fraction of genes, ∼2, 000 out of 19, 000 (2). Thus, the large number of protein-coding genes that can be considered to be understudied represents a real limitation to understanding the functional space, and therefore, it is still not possible to create an unbiased and comprehensive set of annotations.

Relying on EI poses certain problems, the first of which is related to the challenge of the identification of orthology of novel genes (3, 4). Secondly, and in the context of functional inference, it is known that different non-homologous proteins, can perform the same role (5), which has at times led to the consideration of functional analogy as a proxy of orthology (6). Indeed, the ‘orthologue conjecture’ (7) is founded on transcriptomics analyses and it states that orthologous genes tend to retain the same function. Similarly, the “one sequence - one structure - one function” (SSF) paradigm (8) also adopts a central position in the function prediction field, given that orthologues are believed to have similar structures in order to perform the same or a similar function, in line with the aforementioned ‘orthologue conjecture’ (7). However, these assumptions might be violated in some scenarios, as with the same level of sequence divergence the functions of orthologues may differ more significantly than that of paralogues (9), challenging the concept of conjecture.

The second problem is related to the use of sequence similarity as a proxy of homology. Traditional (naïve) methods to transfer functions based on similarity use BLAST (10), reciprocal (11) scores or profile-based models (e.g. hidden Markov models: (12)), with the latter being the most reliable since they are robust statistical models. The rationale is that similar genes may perform similar functions and therefore, we accept transferring functions when sequence similarity exceeds a given threshold (13). These approaches have been used for decades, and they are still in use to annotate regions of proteins and they have been incorporated into the core of most modern methods. More recent approaches include phylogeny-aware criteria (14) and these are becoming standards in the field.

However, all these methods fall short when the sequence similarity cannot be recovered or it is difficult to establish, as is the case of intrinsically disordered proteins (15) or when regions of the proteins exhibit certain properties (complexity, repeats, etc.). This is especially critical in order to understand new and distinct biological questions as genomic and/or transcriptomics information from novel organisms becomes available. Non-model organisms pose a real challenge when predicting function as they may contain genes that perform distinct functions and that are orphan or lineage-specific (16, 17). These genes are not commonly detected or annotated, particularly as the latter is a process that mainly relies on information transfer from small sets of model genes (2).

At the core of the problem is our (lack of) understanding of “function”, which is context-dependent. To circumvent this, the Gene Ontology (GO) Consortium (18) designed a classification scheme based on assigning controlled vocabularies that define functions, and it has become the authoritative source to annotate protein function using distinct ontologies. The ontologies are distributed in direct acyclic graphs (DAGs), wherein the nodes are terms that are connected by links indicating distinct relationships. Ontologies aim to capture the complexity of a function and standardize the terms used to assign functions, and they are divided into the ontologies of Molecular Function (MF), Biological Processes (BP) and Cell Components (CC).

The Critical Assessment of Functional Annotation (CAFA) monitors the state-of-the-art in terms of automatic function prediction, measuring how well computational methods perform (19). CAFA runs as a competition and while two additional contests have been conducted since CAFA3, the results of CAFA4 (in 2020) and CAFA5 have yet to be published (https://www.kaggle.com/competitions/cafa-5-protein-function-prediction). From the most recent competition published, CAFA3, predictions for the BP and MF ontologies have improved slightly over time but overall, term-centric prediction was still found to be challenging.

While computer science technologies were introduced years ago to assist in defining biological functions, until now their impact has been more limited than had been hoped. However, the increasing abundance of data and the technological improvements, both in terms of hardware and software, means that these tools are now gaining momentum. For instance, machine learning (ML) methods were initially employed to predict secondary structure (20, 21) and now, with the technological advances and the vast amounts of data available, this is a blossoming field. This is neatly illustrated by the AlphaFold algorithm (22), which has transformed the prediction of three-dimensional protein structures from sequences. However, in the context of function prediction, a similar breakthrough is still to come. Nevertheless, ML methods are truly gaining protagonism in this area, as witnessed in the last published CAFA3 (19) in which a ML based ensemble method (23) performed better than the rest of the methods across three challenges, unfortunately using a tool that is no longer available. Despite these improvements, these novel methods have not produced a remarkable improvement when it comes to predicting protein function, despite their reach (they are powerful) and the data now available. Beyond the context of CAFA3, popular methods involve the use of convolutional networks (DeepGO, DeepGOPlus), which have also been benchmarked against the CAFA3 datasets (24, 25) in independent publications and shown to perform as well as the top performers.

Orthogonal disciplines offer promising approaches and ML is definitely a vast resource to be explored (26). One such field is that of linguistics, where meaning can be inferred from a text and extracted from how patterns of words are distributed (27), taking into consideration the constraints that are applied to words in order to ensure they have meaning. It is known that the structure and function of a protein also constrains how mutations are selected or become fixed during evolution. Therefore, it may be possible to infer some functional information from sequence patterns. In this context protein function may be predicted through transfer learning (28), whereby proteins are represented as continuous vectors (embeddings) using language models (LMs: e.g. 29). For instance, in Natural Language Processing (NLP) the ELMo model has been trained on an unlabelled text-corpora in order to predict the word most likely to appear next in a sentence given all previous words in the sentence. Subsequently, this model uses a probability distribution for sentences to “learn” certain things about language, such as syntax and semantics. The trained vector representations (embeddings) are thereby contextualized within this context, whereby two identical words could have different embeddings depending on the surrounding words. These concepts have been used to train the analysis of protein sequences at two levels: per-residue (word) to predict secondary structure and intrinsically disordered regions; and per-protein (sentence) to predict subcellular location (28). Consequently, it was concluded that transfer learning can succeed in extracting information from unlabelled sequences to predict location.

In accordance with the above, approaches like GOPredSim (1) train LMs on sequences and they transfer annotations using embedding similarities rather than sequence similarity. The rationale behind these approaches is to predict GO terms through transfer learning of annotations based on the proximity of proteins in the LM embedding space rather than the sequence space. The embedding first used originated from deep learned LMs designed for protein sequences (SeqVec, ProtT5), transferring the knowledge gained from predicting the next amino acid in 33 million protein sequences. The GOPredSim method was benchmarked using the CAFA3 challenge and gold datasets, producing similar results to the ten top CAFA3 performers (1). As these methods are agnostic to evolutionary reliance, they could be useful to annotate the protein functions that challenge the SSF paradigm.

Here we have analysed the capacity of these method’s based on LMs to offer precise and informative functional insights into proteins, as well as to recover functional information from transcriptomic experiments carried out on four representative model organisms: *Saccharomyces cerevisiae, Mus musculus, Drosophila melanogaster,* and *Caenorhabditis elegans*.

## MATERIALS AND METHODS

### Datasets used in this study

*Proteomes.* We collected all the proteins in the reference proteomes at https://www.uniprot.org/ for: *C. elegans* (CAEEL, UP000001940), *D. melanogaster* (DROME, UP000000803), *M. musculus* (MOUSE, UP000000589), and *S. cerevisiae* (YEAST, UP000002311).

*GO annotations.* GO annotations were extracted for the corresponding proteins using GOA (Supplementary Table ST4: https://ftp.ebi.ac.uk/pub/databases/GO/goa/UNIPROT/goa_uniprot_all.gaf.gz). We used all the annotations, including those that are predictions based on homology, since considering only those for which experimental evidence is available would severely limit the coverage.

*Mapping of identifiers.* Entries from the proteomes were mapped to genes through https://www.uniprot.org/id-mapping (Nov 2023, Supplementary Table ST5).

### Transcriptomics studies used in this work

Published data was collected and lists of differentially expressed genes (DEGs) were extracted to perform further enrichment analyses (see below). The genes selected aimed to encompass different strategies and goals for transcriptomics analyses: systemic global changes and temporal serial changes. When possible (according to any differences in the identifiers and versions used), we mapped the specific differentially expressed identifiers to UniProt annotations and to the prediction datasets in each experiment. The datasets used were (Supplementary Table ST9): *S. cerevisiae* transcriptomics data in response to heat shock (30); *C. elegans* classic experimental data that focused on lifespan (31) and on the effects of *Nuo-6* knock-out on the organism; for *D. melanogaster* we selected a temperature-acclimatization transcriptome response (32); for *M. musculus* we selected a pre-B cell differentiation experiment in a model cell line (33).

### Prediction methods used in this work

#### A profile-based method.

This method was used as our internal control, employing the HMMER suite (12) to perform profile-based searches against the Pfam-A database (https://www.ebi.ac.uk/interpro/download/pfam/). Searches were carried out to identify protein domains and in order to detect any GO terms associated with the particular domain. In order to assure that we are recovering homologous domains, we used the independent HMMER reported e-value (i.evalue) since it is more stringent. The results were filtered conservatively by keeping the best domain i.e-value (≤ 0.001), that with the longest alignment coverage that avoids overlapping domains at a minimum alignment coverage of 70%. We selected The GO annotations associated with the protein domains identified were extracted from pfam2go (Supplementary File S1, http://current.geneontology.org/ontology/external2go/pfam2go).

#### Deep Learning (DL) based methods

Two DL methods were selected: DeepGO (24) and the enhanced DeepGOPlus version (25). These methods are DL based approaches that also use EI, as well as protein-protein interaction (PPI) data, in the framework of a Convolutional Neural Network (CNN). For DeepGO, as the code was not available, we used the API provided to retrieve the results (https://deepgo.cbrc.kaust.edu.sa/deepgo/api/create). By contrast, DeepGOPlus was installed locally and the parameters were modified to enable annotations in proteins longer than 1002 residues. As these methods annotate all entries at the root of each category, we only kept those GO terms at a depth >2 from the root of each GO category to avoid annotations at the root.

#### Language Model (LM) based methods.

We used the GOPredSim (1) implementation of two different LMs (SeqVec and ProtT5). To run these methods, protein embeddings are calculated (per protein) for the test dataset. The distances between the protein embedding space were then calculated against the embeddings of a reference dataset (GOA2022, entries with curated GO information), after which a “Transfer Learning’’ approach was applied based on these distances (the smaller the distance, the more likely to transfer the annotations). For the LMs, proteins longer than 5, 000 amino acids were excluded from the predictions since longer lengths are not accepted by the methods (these represent less than 1% of the proteins in our datasets).

In order to avoid overfitting we recompiled the look-up tables, removing the entries of a particular organism prior to running predictions on that particular organism (i.e.: when predicting for *D. melanogaster* any entries or embeddings from this organism were removed from the look-up table, and the same procedure was carried out for the other organisms). This procedure was performed on launching the methods.

All these methods were run on an AMD Ryzen 9 5950X 16-Core Processor CPU with 128GB RAM. Embeddings for both methods were calculated on a NVIDIA GeForce RTX 3090 Ti GPU (24 GB VRAM). The results can be found in Supplementary File F2.

### Estimation of the measures derived from GO terms

#### GO Term precision

Mapping to GO terms was achieved using the GO.db R package (https://bioconductor.org/packages/release/data/annotation/html/GO.db.html) and based on the data provided by GO (https://current.geneontology.org/ontology/go-basic.obo) with a date stamp from the source of 2023-01-01. To check how well the methods predict the exact GO term annotated by UniProt, we devised a 4 type classification to assess the precision of the match by looking at the GO DAG for each organism (Supplementary File S3): “HIT” indicates when the method predicts the exact GO term as it appears in the UniProt annotation; “Close” when the prediction is either a child or a parent of the correct term in the DAG; “related” when the method predicts either an ancestor or descendant in the DAG; and finally, “unrelated” when the predicted terms don’t lie in the branch or that lie beyond any related terms in the DAG. We recorded the precision classification for each method and for all the proteins in the different organisms, computing their frequency.

#### GO Term specificity

The DAG reflects the GO term’s structure and a straightforward way to calculate the specificity of a term is by counting the number of jumps required for a given term to reach the root node. Ideally, that would be most useful if the number of children per branch is similar and the number of children per node is more or less homogenous. In that case, we could assume that each node and each level of the node would be equally informative. However, this is not the case if we have branch A with 4 levels and another branch B with 11 levels, and the nodes of level 3 in both branches are not equally informative.

An alternative way to deal with this situation is to calculate the Information Content (IC) for each node. The IC depends not only on the distance towards the root node but also, on the number of descendants the node has (the greater this number, the less informative it is) and the probability that this term would appear in an annotated set (i.e.: a very specific term should appear less frequently in the dataset of an organism than in the more general dataset).

Using the GoSim package (34), the formula to calculate the IC was:

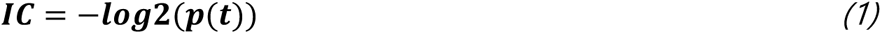

It is defined as the negative logarithm of the probability of observing t, where t is a particular GO term and p is the probability of that term occurring in a dataset, such that p(t) is calculated as:

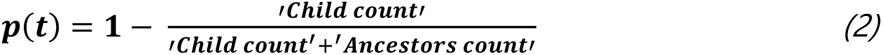

The probability of a term occurring is the frequency with which that particular term or any of its children/ancestors occur in a dataset. GO terms that occur less frequently have a higher IC and are assumed to be more specific. The child and ancestor count refers to the number of proteins that are assigned to that term given a reference dataset, in this case the OrgDb R package (see Supplementary File S4).

#### Similarity among GO terms

The rationale behind assessing semantic similarity (SS) resides on the assumption that genes with similar functions should have a similar annotation vocabulary and a close relationship in the ontology structure. SS among groups of GO terms was calculated using the GoSemSim package (35), employing the Wang method (36) that determines the SS of two GO terms based on both their locations in the GO graph and their relationship to their ancestral terms. This approach enabled quantitative comparisons to be made by first defining the semantic value of a term A as the aggregate contribution of all terms in the “A” DAG to the semantics of term A. Thus, terms closer to term A in the graph contribute more to its semantics.

A protein can be annotated with multiple GO terms and hence, SS among proteins needs must be the aggregate of different SS scores for multiple GO terms associated with these proteins. To combine the SS of our annotations and the gold standards we used the Best-Match Average (BMA) method (https://yulab-smu.top/biomedical-knowledge-mining-book/semantic-similarity-overview.html#bma), which calculates the average of all the maximum similarities (Supplementary File S5).

### Distribution of the scores

For each method, the scores were distributed in quantiles using the R quantile function. This function orders the scores and distributes them into their quantiles, creating 4 groups. In the graphs: the first group will contain all the annotations, regardless of the score; the second group contains the annotations with the first 25% of the annotations removed, those with the lowest scores; in the third, 50% of the annotations were removed and the 50% with the highest scores were retained; and in the last group 75% of the annotations were removed, leaving only the 25% with the highest scores remaining.

### GO term enrichment in transcriptomics experiments

We transferred all the protein annotations to mapped genes. As the methods used provide different scores for similar proteins, when different proteins show distinct scores for the same term and these proteins are associated with the same gene, we kept the highest score for the particular term from the protein pool (Supplementary File S2).

We extracted the sets of DEGs from each experiment (genes of interest, Supplementary Table ST6) and performed GO enrichment tests (see below) on them in all the GO ontologies, using: i) the GO terms extracted from the UniProt annotations; and ii) the GO terms predicted by the methods. Enrichments were performed against all genes from each organism’s genome as the universe (Supplementary Table T7 and as shown in Supplementary File S7).

#### Effect of the scores

As each method has a different scoring system and the scores are not comparable (see results), we distributed the annotations for each method into four groups based on their scores using the R quantile function, a tool that orders the scores and then computes their quantiles (see above).

#### Enrichment tests

we used TopGO (37) <u>10.18129/B9.bioc.topGO</u>, R package version 2.50.0), setting a Fisher’s exact test and using the "elim" algorithm (38) for all the methods in the BP, MF and CC ontologies. We kept the classic Fisher value of p <0.05 to extract significant terms (Supplementary File S6).

#### Semantic similarity (SS)

We calculated the SS among the group of enriched terms from each method relative to the groups of terms from the “annotations” using the GoSemSim package (35), and using the combined BMA metric (as indicated above and see the data in Supplementary Table ST9).

## RESULTS

### Prediction methods increase the annotation coverage that can be achieved in the proteomes of model organisms

Most gene annotations are assigned to the products of genes (proteins) and therefore, here we have focused on the proteomes of four model species (39). From the perspective of function, GO (18) provides annotations in the format of controlled vocabularies that are structured into three DAGs, one for each ontology: BP, MF and CC. However, the annotation status for each organism varies considerably, with *C. elegans* the organism that has fewer annotated proteins and less coverage in all these ontologies (Figure 1a, orange bar).

**Figure 1.**
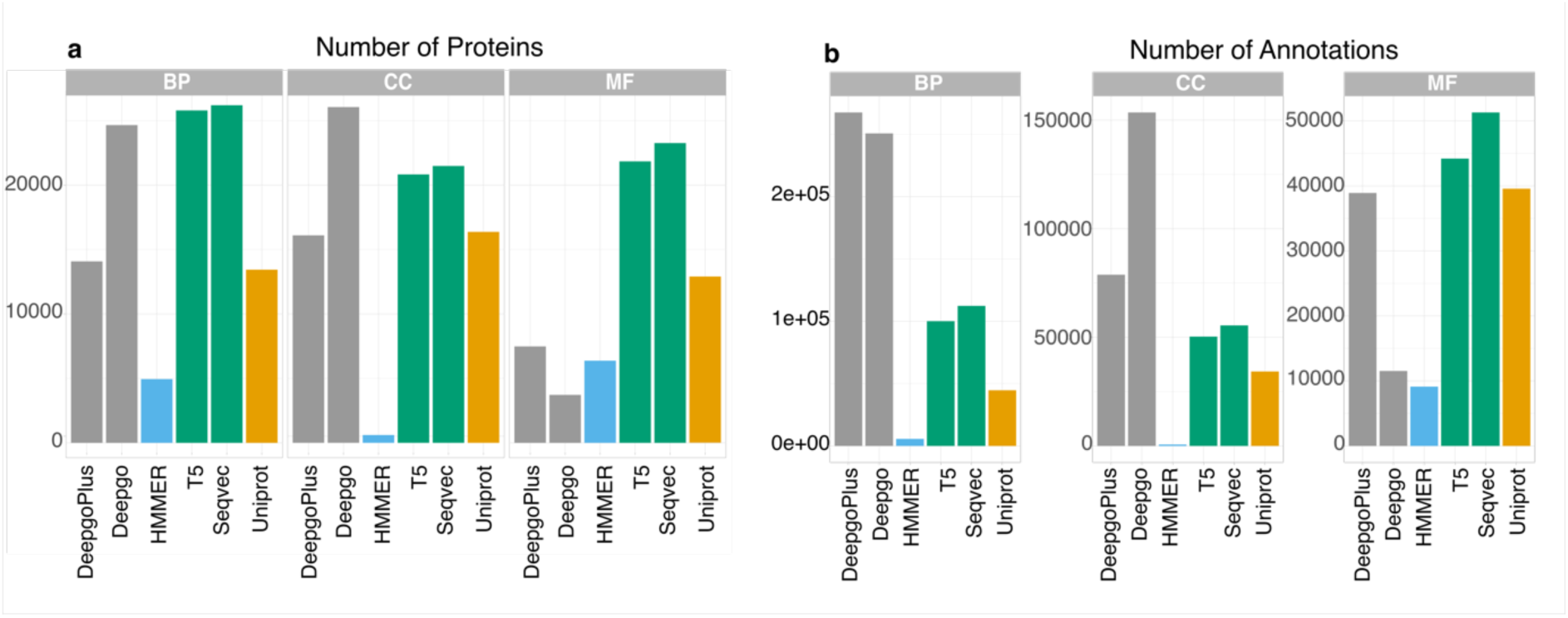
Annotations and predictions in *C. elegans*. **a.** The number of annotations (orange) and predictions (other colours). **b.** Annotated proteins (orange) and proteins with predictions, from a total number of 26, 664 proteins: BP (Biological Process), CC (Cellular Component), MF (Molecular Function). The Language Model (LM) based methods (T5 and SeqVec) are in green, the Deep Learning (DL) based methods (DeepGO and DeepGOPlus) are in grey, the data from a profile-based method (HMMER, our internal control) is in blue, and the annotations extracted from UniProt are in orange. For the remaining species see Supplementary figure SF1).

When drawing up these annotations, despite removing sequences and embeddings from the proteins that correspond to organism targeted, there are still sequences present from highly related species in the training dataset. That is, despite the removal of mouse proteins when targeting that organism, very similar human sequences still remain, which could explain the performance outcomes observed in mouse and fly. Therefore, here we focused on the least annotated species, *C. elegans*, as it is less prone to receive information from similar species. Indeed, in this species ∼72% proteins are annotated (19258 GOA annotated out of 26664 proteins in UniProt, Supplementary Table ST2). Nevertheless, for the remaining species considered, and despite being better annotated, proteome annotation is still to some extent incomplete (Supplementary Figure SF1, Supplementary Table T2, T3, Supplementary file S2). Hence, there would appear to be a non-negligible fraction of proteins that remains unannotated, even in those species that have been studied more extensively (Supplementary Table ST8).

In general, the methods employed here increased the number of proteins with predicted annotations (Figure 1b, Supplementary Figure SF1) and as a general trend, the BP category received more predictions than the other two categories (Figure 1b for *C. elegans,* Supplementary Figure SF1, Supplementary Table ST1). However, the relationship between the number of proteins and the annotations per protein largely depends on the method used. In general, both LM methods were more consistent in terms of the number of proteins with predictions and the median number of predictions per protein (supplementary Table ST1) than the DL methods.

There was wider variation in the values reported with DeepGOPlus, although a similar number of annotated proteins were obtained relative to those achieved with the LMs (see supplementary Table 1), except for in the CC ontology. The MF category was that for which DL methods provide fewest annotations. Then, in *C. elegans* (Figure 1b), for the MF ontology, T5, SeqVec, DeepGOPlus, and DeepGO annotate 21, 839, 23, 262, 3, 720, and 7, 482 proteins, respectively. For CC, they annotate 20, 829, 26, 197, 26, 054, and 16, 100 proteins, respectively. Finally, for BP, they annotate 25, 793, 26, 197, 24, 656, and 14, 080 proteins, respectively.

Regarding the median number of annotations per protein (Supplementary Table ST1), DL methods provide larger numbers per protein than LM. For the MF ontology, T5, SeqVec, DeepGOPlus, and DeepGO produce 2.02, 2.21, 3.1, and 5.2 annotations per protein, respectively. In the CC ontology, they produce 2.42, 2.58, 5.89, and 4.89 annotations per protein, respectively. Finally, for the BP ontology, they produce 3.88, 4.29, 10.17, and 19 annotations per protein, respectively.

We used the profile-based HMMER method as an internal control, a very accurate method but with low coverage that annotates ∼0.29 proteins (Supplementary Table ST2) obtaining about 1 prediction per protein in all ontologies. To estimate how the GO predictions obtained by HMMER (PFAM searches) were related to the predictions generated by the other methods, we identified the GO terms that coincided between HMMER and any other method in *C. elegans* (Supplementary Table ST1). The overlap among the GO annotation terms that coincided between HMMER (PFAM) and any other method was very low, at most 10% in the MF category. Thus, the remaining annotations were further analyzed in terms of their precision and coverage (see below).

While *C. elegans* was the worst annotated of the set of model organisms, it greatly benefits from these predictions in terms of coverage. LMs provided a large number of predictions in all ontologies relative to the available UniProt annotations, except for *S. cerevisiae* CC (Supplementary Figure SF1). Interestingly, of the DLs, DeepGOPlus performed worse than HMMER in terms of retrieving predictions for the MF category, which is itself a low coverage method, although DeepGOPlus was the method that provided more annotations for CC. It is interesting to note that the LMs annotated more proteins than the DLs in MF and BP, except for in *D. melanogaster*. Thus, while the LMs increased the coverage of *C. elegans* in all ontologies, this trend was only evident in the BP for *D. melanogaster* and *M. musculus*. These results may reflect the focus of different scientific communities on particular genes or gene families, with different groups of genes having been studied more intensely, while others remain relatively overlooked (**2**). Overall, the LMs (GOPredSim-T5 and GOPredSim-Seqvec) provided a larger and more consistent coverage of the predicted functions for proteins across organisms, regardless of their annotation stage, making them promising tools to address functional studies in non-model organisms as already indicated in the animal phyla (40).

### Language Model predictions are more precise than those from Deep Learning methods

The different methods assessed produced a large coverage of predictions, yet how precise are they? To address this question, we estimated the ability of the methods to guess the exact GO terms assigned to proteins (as per the GOA2uniprot annotation) using a scheme based on the localisation of the prediction in the DAG’s relative to the real annotation (Figure 2a). Despite the large number of predictions, BP was the worst predicted category in *C. elegans*, while MF was the best (Figure 2b). In terms of precision, the same trend was observed for the remaining species (Supplementary Figure SF2). This observation was expected since BP is a difficult category to predict without also using text mining and/or network information. As expected, HMMER (our base-control) is very precise at the expense of a very low coverage. When annotations from HMMER were removed, a substantial fraction of HITs and related functions were still recovered (Figure 2c-d). Hence, LMs, and GOPredSim-T5 in particular, provided a wider annotation range with better precision.

**Figure 2.**
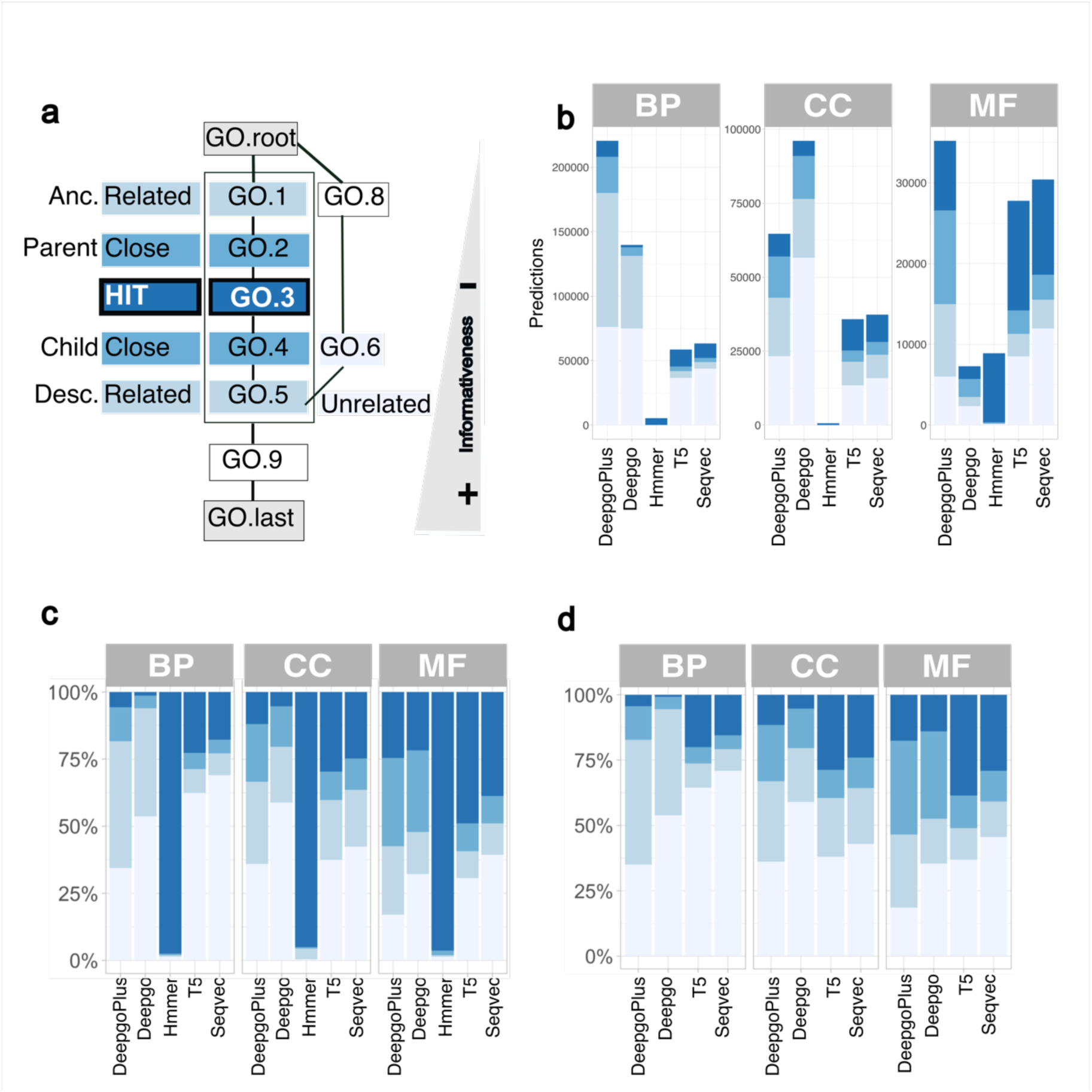
Precision achieved by the different methods in *C. elegans.* **a. Definition of precision** in the context of an hypothetical DAG, where “GO.root” is the root of the category (e.g., BP) and “GO.last” is the most specific term: HIT (darkest blue) matches the term; “Close” is either a parent or child; “Related” includes any descendant/ancestor; and there are “Unrelated” predictions. The informativeness also depends on the position of the predicted term in the graph and it is higher close to the end. **b. Number of predictions** (Y-axis) per method (X-axis) or per category *for C. elegans*, according to the “precision type”. The results for the other species can be found in Supplementary figure SF2, which exhibit similar trends. **c**. **Proportion of the predictions according to precision**. The numbers indicate the type of predictions. **d. Proportion of the predictions according to precision once the GO terms by HMMER were removed.** The numbers indicate the type of predictions one the coincidental annotations are removed (See Supplementary Table ST1). The “precision type” follows the colour scheme of panel ‘a’.

### Language Models are more informative than Deep Learning methods

Once the coverage and precision had been established, we estimated how informative, in terms of detailed annotation, these predictions were. One way to look at the information provided by the terms was to identify the predicted term localisation within the DAG. The closer to the root (less detailed) a term is (Figure 2a), the more general it is and thus, the less informative it is. To quantify this facet we calculated the IC of the predictions retrieved by the different methods, whereby less informative terms (closer to the root) were ascribed lower IC values (Figure 3a). Our results from *C. elegans* indicated that overall, LMs were the most informative methods in the BP and MF ontologies, providing more specific information. Interestingly, HMMER, the baseline method, performed better for the CC category in all species.

**Figure 3.**
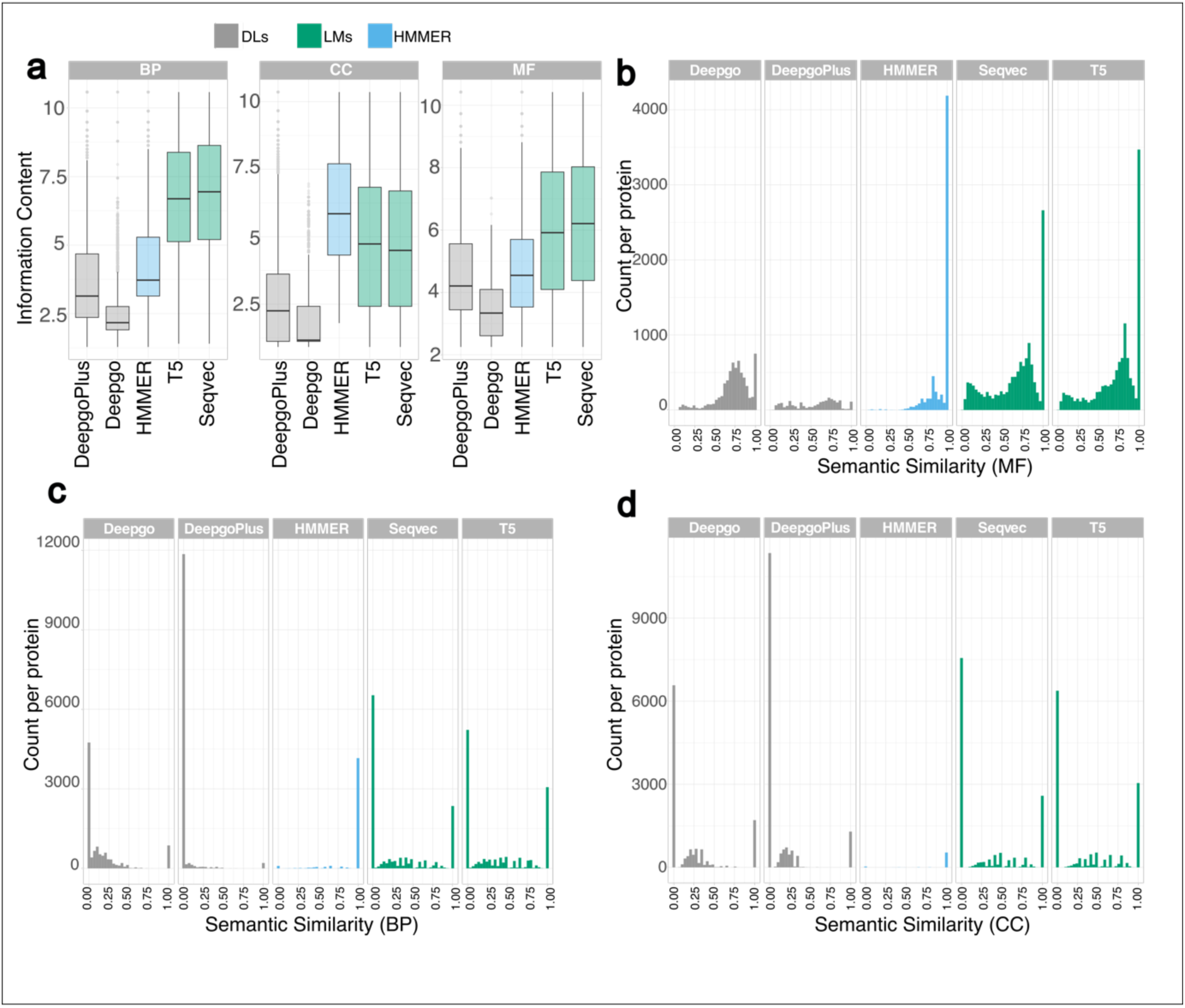
Information extraction in *C. elegans*. **a. Information content (IC).** The IC (Y-axis) values for each method across the ontologies: the larger the number, the more informative (more specific terms). **b.** The Semantic Similarity per protein (Y-axis) represents the protein counts and the X-axis indicates the SS value where 1 indicates that the annotations and predictions are identical for MF, **c.** for BP, and **d.** for CC. The results for the other species can be found in Supplementary figure SF3, which exhibit similar trends.

In addition to the IC, we also estimated the SS among the predicted GO terms relative to the GO terms annotated per protein, where values close to 1 indicate a high similarity among the group of terms (Figure 3b-d). We found that MF was the best predicted category (Figure 3b), that which provided more similar terms, while the BP and CC categories (Figure 3c-d) provided many unrelated predictions per protein relative to the standard annotations. The same trend was observed for the other species (Supplementary Figure SF3).

### Language Models provide a better precision and coverage per protein than Deep Learning methods

In order to check the performance of these methods in terms of protein coverage, we must consider two different but related aspects: the number of annotations that are correctly predicted by a method with a given precision (Figure 4a left); and how many of the predictions coincide within the per protein space of annotations (Figure 4a, right), i.e. does the method predicting everything that can be predicted? For instance, HMMER predicts ∼5000 *C. elegans* proteins to have a GO term match as a HIT in the BP ontology (the predicted GO terms are UniProt matches), although it did not predict all the GO terms that could be predicted. In other words, while most HMMER predictions for a protein were perfectly guessed (Figure 4b, left), this method missed a substantial number of predictions (Figure 4b, right). Notably, in terms of precision LMs outperformed DLs at the highest level of precision (“guessed”, upper level). The same trends were also observed in the remaining species (Supplementary Figure SF4).

**Figure 4.**
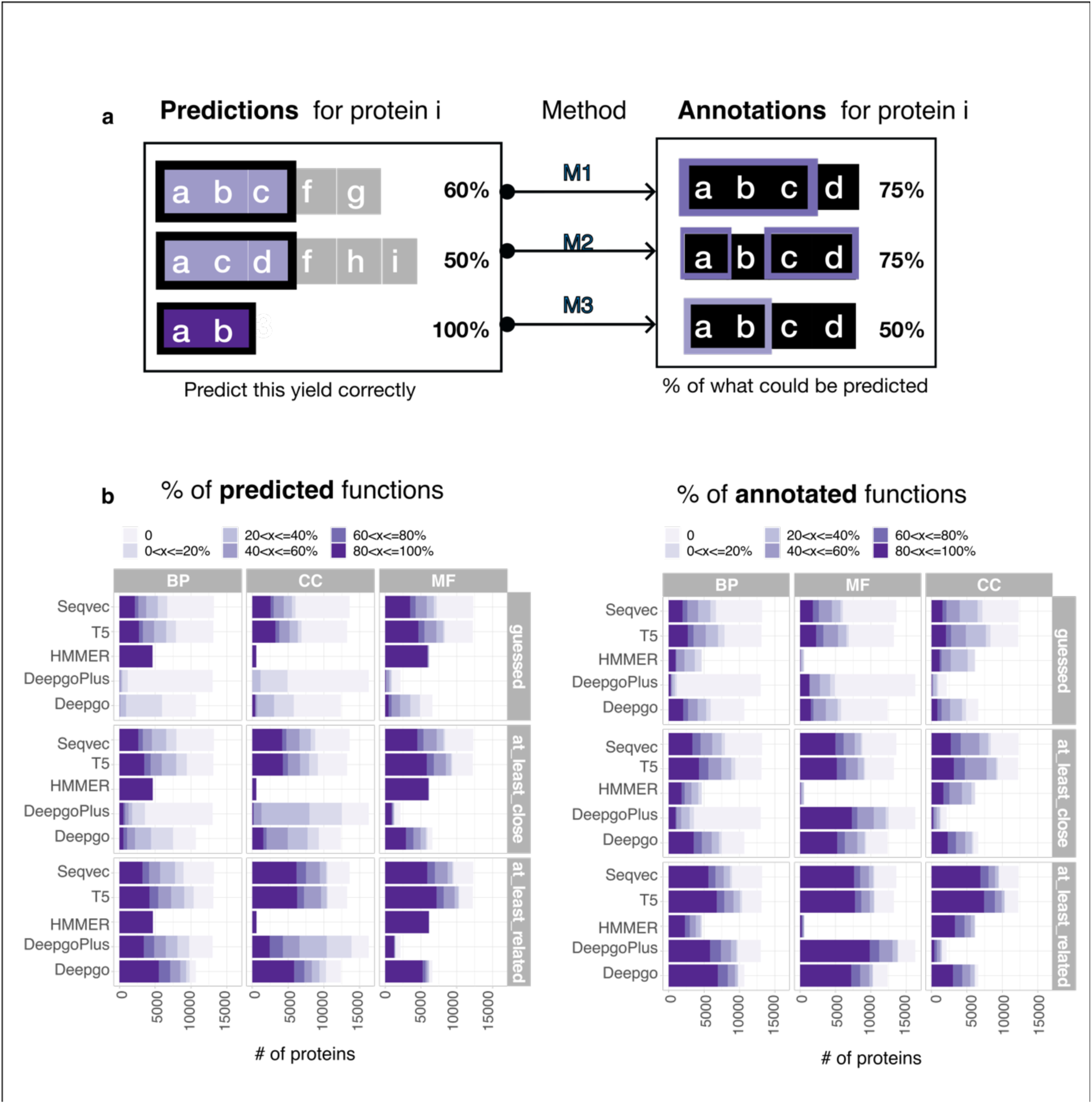
Per protein coverage of the methods. **a.** From the entire prediction space, the left side represents how many are correct predictions (annotated), while the right side represents the fraction of correct predictions out of what could be predicted from the annotation space. **b. The results for *C. elegans*:** Left, the proportion of predicted functions across ontologies; right, the proportion of annotated functions. The colours indicate the ranges in percentages. The results for the other species can be found in Supplementary figure SF4, which exhibit similar trends.

### The scores have an effect on information retrieval and coverage

With the exception of HMMER, where selection was based on i.evalue and alignment length (see Materials and Methods), each method analysed here produced different scores. The scores were not readily comparable since DLs do not fully cover the range of 0-1 (Figure 5a): the DeepGO method (API version) filters the scores beforehand, producing scores from >0.3-1; while the DeepGOPlus scores vary from 0.1-0.5. We then distributed the predictions for each method into four groups according to the quantiles of the scores (see Methods), assuming that the last group will concentrate the predictions with the highest scores, in principle the most trusted predictions.

**Figure 5.**
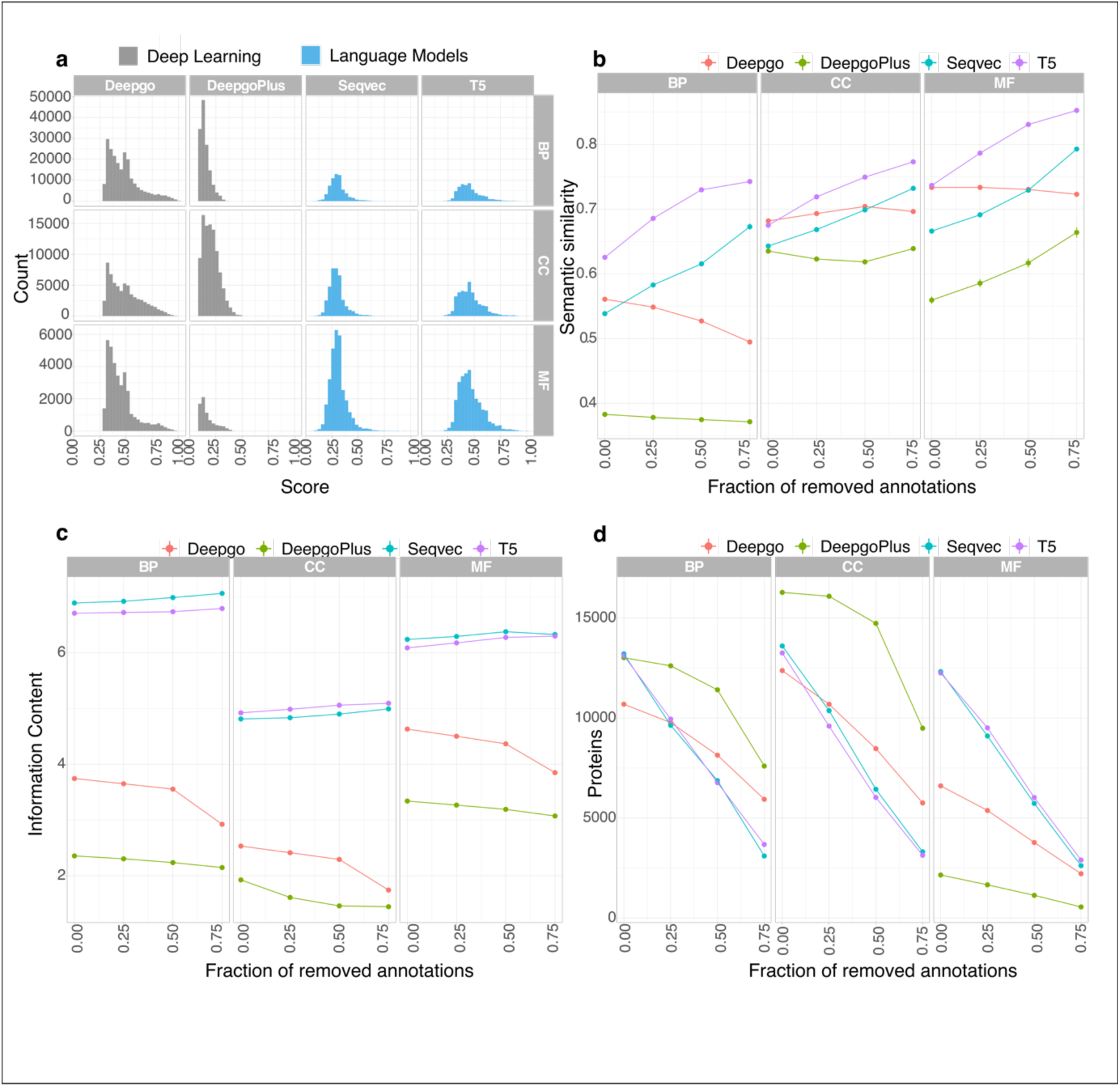
Score effects in *C. elegans*. **a. Score distribution of methods.** The distributions of the scores are not comparable between LMs (language models) and DLs (Deep Learning methods). **b.** The effect of the score on the Semantic similarity (SS). The Y-axis represent the SS values (SS >0.7 are considered good starting values), while the X-axis represents groups of annotations distributed according to their increasing scores, where: 0.00 includes all the annotations regardless the score; 0.25 includes annotations where 25% of the lowest scoring annotations are removed; in the 0.50 group 50% of the lowest scoring annotations are removed; and in the 0.75 group, 75% of annotations were removed leaving only the annotations scoring highest in this quantile. **c.** The score’s effect on IC (Y-axis) and the X-axis is the same as in panel a. **d.** The score’s effect on protein coverage: the Y-axis is the number of proteins harboring annotations at these scores; the X-axis is the same as in panel a. The results for the other species can be found in Supplementary figure SF5, which exhibit similar trends. Each method studied is depicted by a different colored line.

#### Effect on Semantic similarity

To assess whether the scoring reflects the recovery of “true” functions, the SS was used as a proxy to measure the similarity among the groups of predictions and annotations, calculating how similar their GO terms are for each protein. As expected, and in line with the results regarding precision, both LMs tended to provide the higher SS values in the last group across ontologies, in which 75% of the annotations had been removed (Figure 5b *C. elegans* and in Supplementary figure SF5 the same trend was evident for the remaining organisms). GOPredSim-T5 is the method that provided the best SS across ontologies and among species, and the SS values increased relative to the increasing scores. The same trend was observed with DeepGOPlus for the MF ontology although it did not reach values >0.7.

Interestingly, when all the scores were considered the SS for BP in *C. elegans* was very low. DLs exhibited a very heterogeneous trends across species and across ontologies, with DeepGOPlus the method that provided the lowest SS in all the ontologies, with a variable effect depending on the species and/or ontology. LMs predicted the BP and CC ontologies worse in *D. melanogaster* and *C. elegans*, while *M. musculus* was the species in which all ontologies were best predicted (Supplementary figure SF5, lowest panel).

#### The effect of scores on the Information Content

Another way to look at the “reach” of information is to estimate the detail of the predictions, with the IC providing a measure of the specificity of the terms (Figure 2a). To address the effect of the scores on the IC (Figure 5c), we plotted the IC according to the the score distribution as indicated above. For LMs, IC values did not show any particular trend, whereas they decreased for DLs. This suggests that while the IC of LMs is not affected by the score, the “best” scores for DLs will produce less detailed information (Supplementary figure SF5 and Supplementary File S3).

#### Effect of the scores on protein coverage

Higher scores should technically provide more accurate predictions as a higher SS implies that GO terms are more similar, irrespective of the specificity of the information reflected by the IC (Figure 5b, c). We expected to observe a substantial drop in proteins assigned high scores for the annotations/predictions based on the distributions. Indeed, there was a significant drop in proteins as the score increases for *C. elegans* (Figure 5d), a trend that persisted in the remaining organisms (Supplementary figure SF5). Accordingly, this could serve as a guide to select scores for different purposes. For instance, if we want to keep the largest coverage at a feasible SS in the GOPredSim-T5 (i.e.: SS >0.7), then predictions excluding the first group (the removal of 25%) will suffice for all ontologies as they still contain a large proportion of the proteins. By contrast, if our aim was to detect the highest SS, then we should focus on proteins in the last group in which the fraction of removed annotations is 0.75.

### Language models recover functional information from transcriptomics experiments

One key approach in functional genomics is to perform GO enrichment experiments under different conditions to identify functions that are specifically enriched or depleted. Assuming that the functional information from transcriptomics experiments reflects the “true” function, we estimated how the predictions recapitulated the annotations in GO enrichment experiments by performing GO enrichment tests on published DEGs. We compared the GO enrichments from predictions against those from annotations by estimating the SS of the results. Protein predictions were first transferred to their cognate genes and when redundant predictions from different proteins pointed to the same gene, we retained the highest score. We then again distributed the predictions into groups according to score’s quantile distribution for each GO ontology, allocating the proteins harboring the scores to the corresponding group (see Methods). The SS values of the predictions and annotations for DEGs were then represented across ontologies for the *C. elegans Nuo-6* KO (Figure 6 down regulated in ‘a’ and ‘c’; up-regulated in ‘c’ and ‘d’: (31)). For LMs, the SS barely decreases across quantiles (Figure 6a, b and Supplementary Figure 6 for the remaining species), while the number of genes harboring significant (predicted or annotated) terms decreased largely as an effect of the score (Figure 6c, d and Supplementary Figure SF6). In other words, at a higher SS there are fewer proteins harboring very similar GO terms. Overall, and in line with previous comparisons, the SS is higher with the LMs for all the ontologies. In this case, LMs can recover the “true” function for the down regulated genes at high SS (>0.8), while for the up-regulated fraction of genes MF (as seen previously, Figure 3b) and CC were the best predicted categories, BP was the worst. As expected, the number of genes annotated drops as the score increases (X-axis, Figure 6c, d). Hence, while LMs provide the highest SSs (Supplementary Table ST9), DL methods can be outperformed even by the classic HMMER.

**Figure 6.**
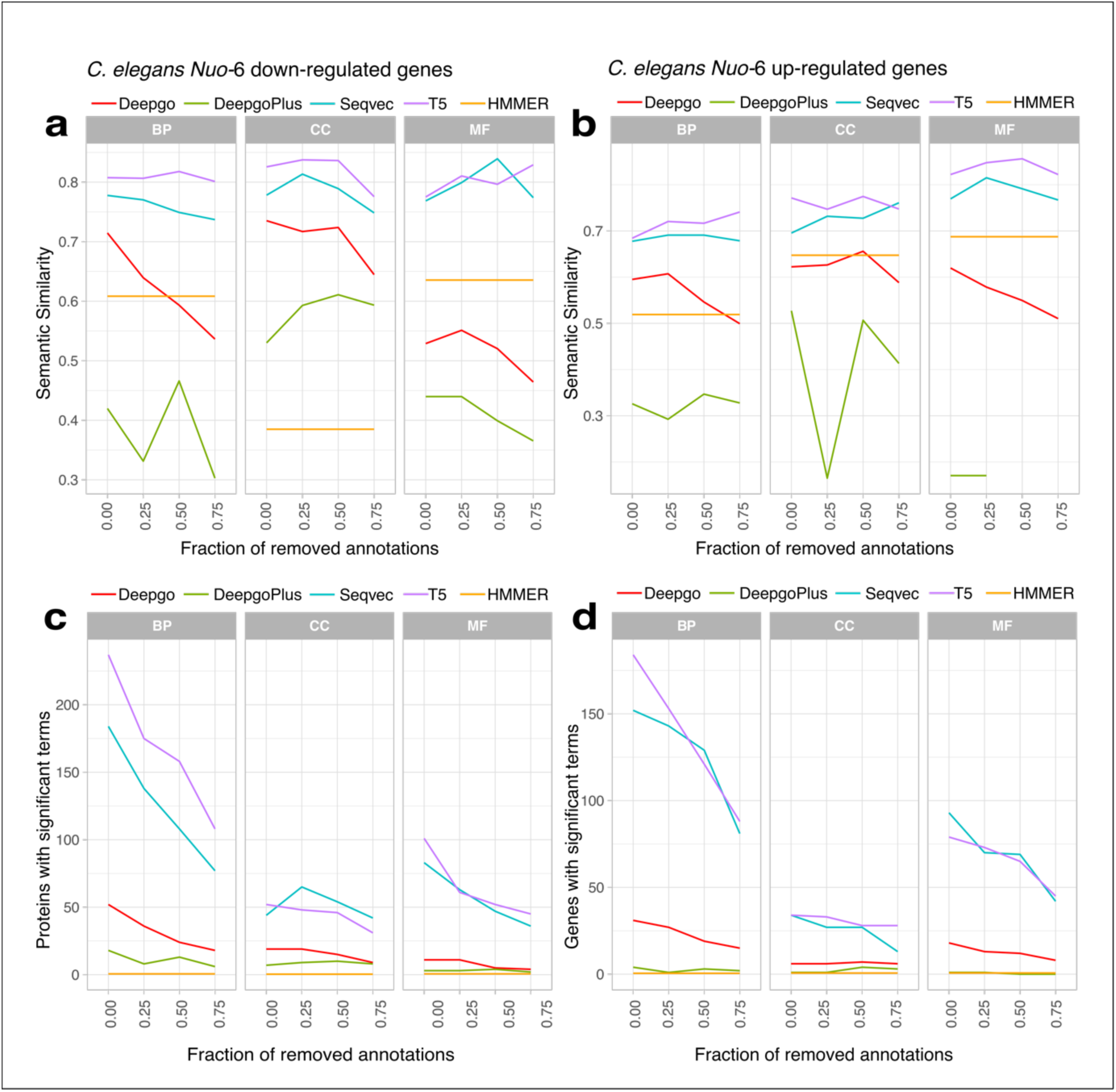
Recovery of functional information for *C. elegans*. Semantic similarity (SS) between GO terms from the TopGO enrichment experiments, the GO terms from annotations in UniProt are compared to the GO from predictions: upper panels, the SS across the score quantiles; lower panels, the number of proteins harbouring significant terms across quantiles. The results for the other transcriptomics experiments can be found in Supplementary figure SF6, which exhibit similar trends.

Overall, LMs can recover “true” functional information regardless of the score and protein depletion, whereas DLs do not show the same trend. Thus, LMs appear to be suitable for full genome annotation if the aim is to perform further enrichment comparisons.

### LM methods complement existing proteome annotation

We found unannotated proteins with predictions (missing annotations in at least one ontology) and the methods that contributed most to these were the LMs, particularly when applied to the BP and MF ontologies, whereas DLs annotated more proteins in the CC. It is interesting to note that HMMER still provides annotations to proteins that in principle should have been annotated by automatic methods, as this method is at the core of most automated predictions. Given that the scores of the methods are not comparable (see above), we selected examples at the top end of each scoring range (Supplementary Table T8). In *S. cerevisiae*, there were still 1, 012 proteins partially annotated by UniProt (missing annotations in at least one ontology) and the methods studied complemented these annotations. Nevertheless, 3 proteins had no annotations for any category in UniProt but all with evidence at the protein level, with P47129 (Assembly-complementing factor 4) receiving annotations in all ontologies using the methods tested here, and with GO terms indicating a role in autophagy and protein transport (GOPredSim-T5), as well as DNA repair (GOPredSim-Seqvec). In *C. elegans*, 9, 158 proteins were partially annotated in UniProt and they missed annotations in at least one category, with 86 of these not annotated at all by methods that provide a wealth of annotations for exploration. For instance, the Glutathione S-Transferase gst-39 (Q9NAB0) receives annotations from GOPredSim-T5 in BP (glutathione metabolic process, prostaglandin biosynthetic process) and CC (intracellular membrane-bounded organelle). Another interesting protein is edg-1 (Q17436), which received annotations from GOPredSim-T5 in BP (developmental-related processes) and in CC (nucleus). While 6, 130 proteins were partially annotated In *D. melanogaster*, 35 had no annotations. For example, the “Common Dpr-interacting protein” (Q9VDD5) is a LRR protein annotated by GOPredSim-T5 in BP (germ cell development, moulting cycle, collagen and cuticulin-based cuticle, by GOPredSim-T5), in MF (RNA helicase binding) and CC (P-granule, plasma membrane). LRR proteins are known to be involved in immune responses and promoting protein interactions (41). This protein does not have a clear human ortholog. The UPF0184 protein CG14818 (O97420) does not have a clear homologue from better annotated species, and also receives interesting annotations in BP (cellular heat acclimation, regulation of transcription by RNA polymerase II). Similarly, the AT27166p protein (Q8T3W6, the “hale gene”) is involved in viral release and proteolysis according to GOPredSim-T5 (BP) and has no clear homologue with protein evidence in better annotated species.

In the particular case of mouse, 16, 315 mouse proteins were partially annotated, with 78 unannotated proteins. These assignations may be largely influenced by the presence of embeddings extracted from very similar proteins from sister species (i.e. *H. sapiens*), where annotations produced by the methods recapitulate those that could be easily transferred from homologous proteins. For instance, the TBK1 binding protein 1 (Z4YKN2) was predicted by GOPredSim-T5 and GOPredSim-SeqVec to participate in I-kappaB kinase/NF-kappaB signalling and in innate immune response in the BP. Similar terms were found in the human homologue, that has been implicated in mediating both tumor growth and immunosuppression (42). However, other methods indicated more general terms like the regulation of cellular process (DeepGO). Additional examples are the Netrin G1 protein (F8WJ48), that received predictions in BP (substrate adhesion-dependent cell spreading, modulation of chemical synaptic transmission, nervous system development) and MF (cell-cell adhesion mediator activity). The BCL2/adenovirus E1B 19 kDa-interacting protein 2 (Q91VL0) was predicted to be involved in signalling and calcium binding: MF (GTPase activator activity, calcium ion binding) and BP (apoptotic process, regulation of catalytic activity).

Finally, we calculated the distributions of scores across our precision categories in the three ontologies (Supplementary Figure S7-9). We found that in mice, for GOPredSim-T5, the “HIT” type concentrated the highest scores across categories, a trend that was not observed in the remaining species nor in the other methods.

## DISCUSSION

Current genomic sequencing projects are producing large amounts of data and in recent years, this is especially true for non-model organisms. However, the experimental determination of protein function cannot keep up with the availability of sequences and hence, the assignation of functions based on experimental testing represents only a small fraction of the sequence universe, compromising any benchmarking approach. In our case, using these approaches alone would definitely compromise coverage. Thus, for non-standard organisms most predictions are homology based, suitable alternatives to increase the functional space. However, traditional homology-based methods may fall short in certain contexts, as there may be a large divergence that is difficult to capture through evolutionary signals but that may be detected using structure-based methods, even though these functions are not necessarily transferable. In addition, similarity cannot be used as a proxy of evolutionary relatedness in lineage or taxon-specific genes, and this may be difficult to implement when sequences have particular features like repeats or low complexity regions. Promising computer sciences approaches have been employed for protein annotation, although their practical use is hindered by the trade-off between annotation precision and coverage, and most of the time by their usability, since large queries are not suitable for web-based solutions.

In terms of precision, the approach adopted here aimed to mimic the natural variance of expert procedures to annotate proteins at different levels of detail. For instance, the GO:0016002 term (sulphite reductase activity) in the MF GO graph has a child term GO:0004783 (sulphite reductase -NADPH), and a parent term GO:0016667 (oxidoreductase activity). These three terms point to a different level of detail in the information and hence, a perfect method should ideally match the annotation (defined in this work as a “HIT”). For *C. elegans*, the HIT GO:0016002 is guessed by GOPredSim-Seqvec, although the more specific child GO:0004783 is not guessed by any method and the more general parent GO:0016667 is predicted by Deepgo (Supplementary file 2). Since not all methods annotate at the same level of detail (as human curators do), it is fair to assume that methods perform as well (or as bad) as humans, yet it appears that LMs produce more precise predictions than DLs. Another important issue is whether or not more detailed information provides better biological insights, although this is an issue beyond the scope of this work as it depends on the specific biological question that is being addressed.

Nonetheless, any prediction method would be expected to provide a scoring threshold that enables users to either select or discard predictions. Scores are values that usually range from 0-1 (normally derived from a particular scoring function), and the thresholds for their usage should be recommended by developers. In the methods tested here, it is not obvious how the scores are calculated from the publications, with partial explanations or no clear recommendations about score usage (with the exception of GOPredSim for SeqVec that provides scores for each category). Although the developers benchmark their methods against dedicated functional datasets (e.g., the standard CAFA3: (19)), to the best of our knowledge there is no comprehensive benchmark at an organism level that includes the recovery of functional information. We also analysed the effect of the scoring system for each method on each species, both at the SS and IC levels. We found that the LMs produce more or less the same range of annotations, the same number of proteins annotated and a similar number of annotations per protein, indicating certain robustness. However, both DLs methods performed quite distinctly, with DeepGOPlus performing worse when compared to its predecessor.

When we analysed the score distributions for the terms at our established precision types (Supplementary Figures 7-9), we found that the only species that accumulated higher scores for the guessed (“HIT”) terms was *M. musculus*, and only for GOPredSim-T5. This could reflect the influence of those annotations from sister species (i.e., *H. sapiens*). However, this trend was not observed for the other species nor the other methods. This indicates that for these species, it is likely to find ’true’ annotations in the ’unrelated’ class. However, since the only way to prove that these unrelated functions are actually ’true’ is through experimentation, any conclusion regarding the reach of the unrelated class would remain purely speculative.

Another relevant issue is related to the informativeness of the predicted terms, specifically in terms of SS with real annotations and the location of the predictions in the DAGs (measured as IC). For LMs, the IC barely changes with the scores and it is relatively high at the lowest scores. By contrast, an increase in the DL scores is associated with a decrease in the IC values, indicating that for these methods higher scores are related to more general terms. A reliable scoring scheme would be expected to provide a better SS at a higher score (i.e.: similar GO terms between predictions and annotations) and while this holds true for LMs, the opposite trend was observed for DLs where the higher scores were most dissimilar.

In terms of the ontologies, they were not all equally well predicted by the methods tested. For instance, the BP ontology is that which receives most annotations but it is also that which is more difficult to predict in the least accurately annotated species (*C. elegans* and *D. melanogaster*), as witnessed by the SS. However, GoPredsim-T5 did recover a SS above 0.7 in the higher score (from the group in which 50% of the annotations had been removed).

Finally, we analysed whether these methods recapitulate the extraction of functional information from transcriptomic experiments. LM methods are capable of retrieving similar functions (measured as SS >0.7) in conservative settings, and of attaining higher scores when the proteins bearing significant terms are largely depleted. Our combined results indicate that LMs provide a fair coverage with a good SS, providing more specific information, whereas DL methods offer broader coverage but they yield more general annotations which are less informative. This trade-off highlights the need to consider precision versus coverage in prediction annotation methods.

Our study demonstrates that LMs may be particularly useful to examine poorly annotated genomes, as exemplified here in *C. elegans*, both at the genome level and in terms of the functional retrieval of DEGs. We also performed largescale testing of LMs, in particular applying the GOPredSim-T5 annotation of 2.4 million to genes in various non-model organisms in the animal Phyla, where we found interesting biological insights (see 40).

At a single species level, we tested these methods on transcriptomics data of non-model organisms (unpublished results). In these settings, we recommend to first maintain the standard annotations assigned by the reference repository (e.g., UniProt, based on experimental evidence), since they are probably derived from homology-based methods. The increase in the “annotation” coverage with predicted annotations from LM is good, ideally generated by language models (GOPredSim-T5), while being informative based on the IC and SS. Finally, to increase the background annotations, we could complement the remaining unannotated proteins with DL predictions at any score, since they do provide very general terms.

To summarise, Language Models are suitable methods for large annotation efforts when homology-based does not provide information, and they can recover “functional” information from transcriptomics experiments. Incorporation of embeddings encoded from structural information are approaches deserving further exploration in the function assignation context.

## DATA AVAILABILITY

The results, code and supplementary data files are available at Zenodo: https://zenodo.org/records/10993618

## FUNDING

This work was supported by grants from the Ministerio de Ciencia e Innovación: (MCIN/AEI/10.13039/501100011033 [(PID2021-127503OB-I00) to A.M.R. and I.B.; (PID2019-108824GA-I00) to R.F.; (RYC2017-22492) to R.F.; (MDM-2016-0687) to I.C., P.M., A.R.]. In addition, we acknowledge the funding of: the LifeHUB/CSIC research network (PIE-202120E047) to G.I.M.R., A.M.R., and R.F.; the European Research Council (grant agreement no. 948281) to R.F; the Human Frontier Science Program (RGY0056/2022) to R.F.; and the Secretaria d’Universitats i Recerca del Departament d’Economia i Coneixement de la Generalitat de Catalunya (AGAUR 2021-SGR00420) to G.I.M.R. and AGAUR 2021-SGR00420 to R.F.). Funding for open access charges were provided by the Ministerio de Ciencia e Innovación: MCIN/AEI/ 10.13039/501100011033.

## CONFLICTS OF INTEREST

None to declare

## Notes

https://github.com/CBBIO/func-lm

### Competing Interest Statement

The authors have declared no competing interest.

### Summary of Updates

New figures to describe the scores distributions and also more comprehensive explanations regarding the results of the analyses.

