## Supplementary for "Decoding functional proteome information in model organisms using protein language models"

### SUPPLEMENTARY MATERIALS AVAILABILITY

Results and supplementary data are available at Zenodo as "FuncProtDecode": https://zenodo.org/records/10659694

### SUPPLEMENTARY FILES

**Supplementary File S1**. HMMER results for the selected organisms per protein.

**Supplementary File S2.** DeepGO, DeepGOPlus, GOPredSim-T5 and GOPredSim-Seqvec prediction outputs and scores per protein for the selected organisms.

**Supplementary File S3.** Precision assignments to GO terms

**Supplementary File S4**. Information content (IC) per term distributed by quantiles for the selected organisms.

**Supplementary File S5.** Semantic similarity (SS) per term distributed by quantiles for the selected organisms.

**Supplementary File S6**. Fisher tests and genes in the transcriptomics TopGO enrichments.

**SUPPLEMENTARY TABLES:** presented as an excel file, where each table is an independent spreadsheet.

**Supplementary Table ST1.** Summary of the annotations: DG, DeepGO; DGP, DeepGOPlus; HM, HMMER; SV, SeqVec; T5, ProtT5; Cel, *C. elegans*; Dme, *D. melanogaster*, Mus, *M. musculus*; Sce, *S. cerevisiae*; and GOA, UniProt annotations.

**Supplementary Table ST2.** Protein and Gene mapped fraction per method. Summary of annotations: DG, DeepGO; DGP, DeepGOPlus; HM, HMMER; SV, SeqVec; T5, ProtT5; Cel, *C. elegans*; Dme, *D. melanogaster*, Mus, *M. musculus*; and Sce, *S. cerevisiae*.

**Supplementary Table ST3.** Gene (mapped from proteins) annotation ratio per method and category.

**Supplementary Table ST4.** UniProt Annotations from proteomes extracted from GOA2Uniprot.

**Supplementary Table ST5**. Mapping proteins to genes. The proteome entries were mapped to genes through https://www.uniprot.org/id-mapping (Nov, 2023).

**Supplementary Table ST6.** Differentially expressed genes from transcriptomics analyses. Genes identified as differentially expressed in published transcriptomic analyses to estimate the retrieval of functional information.

**Supplementary Table ST7.** Whole genes in the genomes. All the genes used in the experiments as a universe in the TopGO enrichment analyses.

**Supplementary Table ST8**. Proteins annotated by methods and not annotated by UniProt in at least one GO category. GO terms predicted in proteins but not annotated by UniProt.

**Supplementary Table ST9.** Semantic similarity (SS) between annotations and predictions in transcriptomics experiments. The SS values and the corresponding number of proteins harbouring significant terms as a score quantile distribution.

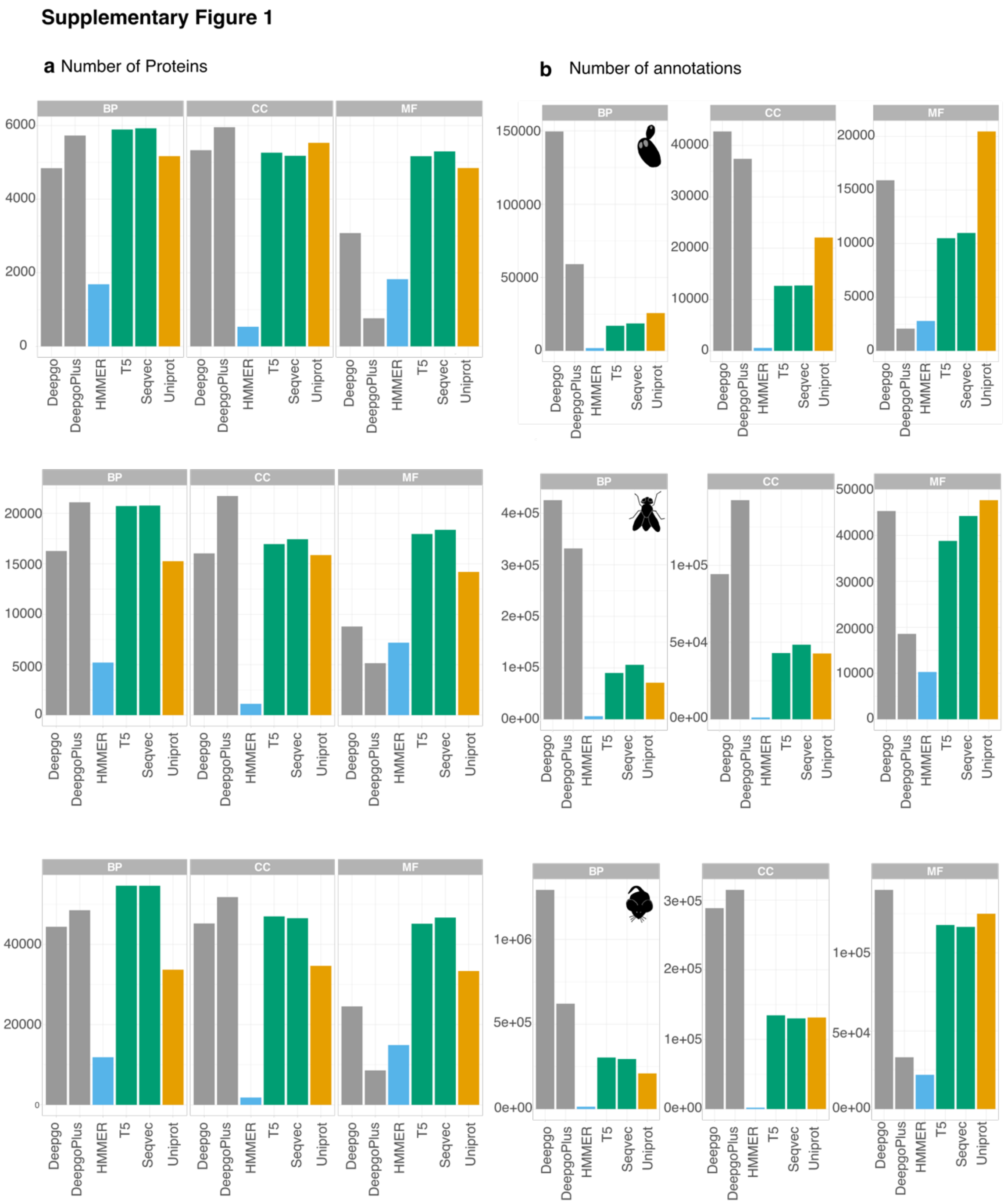


### SUPPLEMENTARY FIGURES

**Supplementary Figure SF1.** Annotations and predictions. **a.** Number of annotations (orange) and predictions (other colours), **b.** Proteins annotated (orange) and proteins with predictions: BP (Biological Process), CC (Cellular Component), MF (Molecular Function). The Language Model (LM) based methods are in green (T5 and SeqVec), the Deep Learning (DL) based methods (DeepGO and DeepGOPlus) are in grey, a profile-based method (HMMER, our internal control) is in blue, and the annotations extracted from UniProt are in orange.

**Supplementary Figure 2**

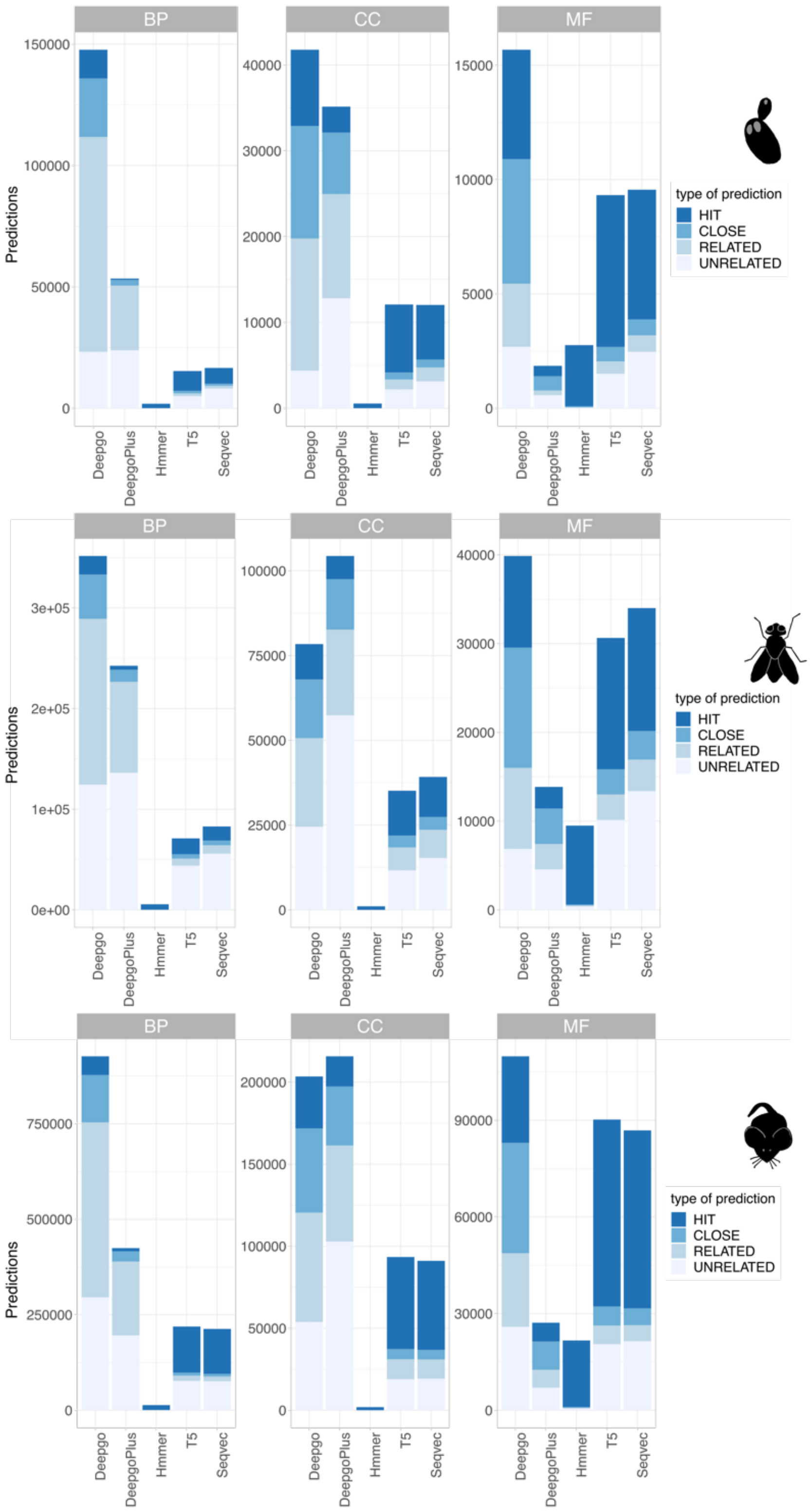


**Supplementary Figure SF2**. **Precision achieved by the methods**. The number of predictions (Y-axis) per method (X-axis) and per category according to their "precision" and following the colour scheme in Figure 2a.

**Supplementary Figure 3**

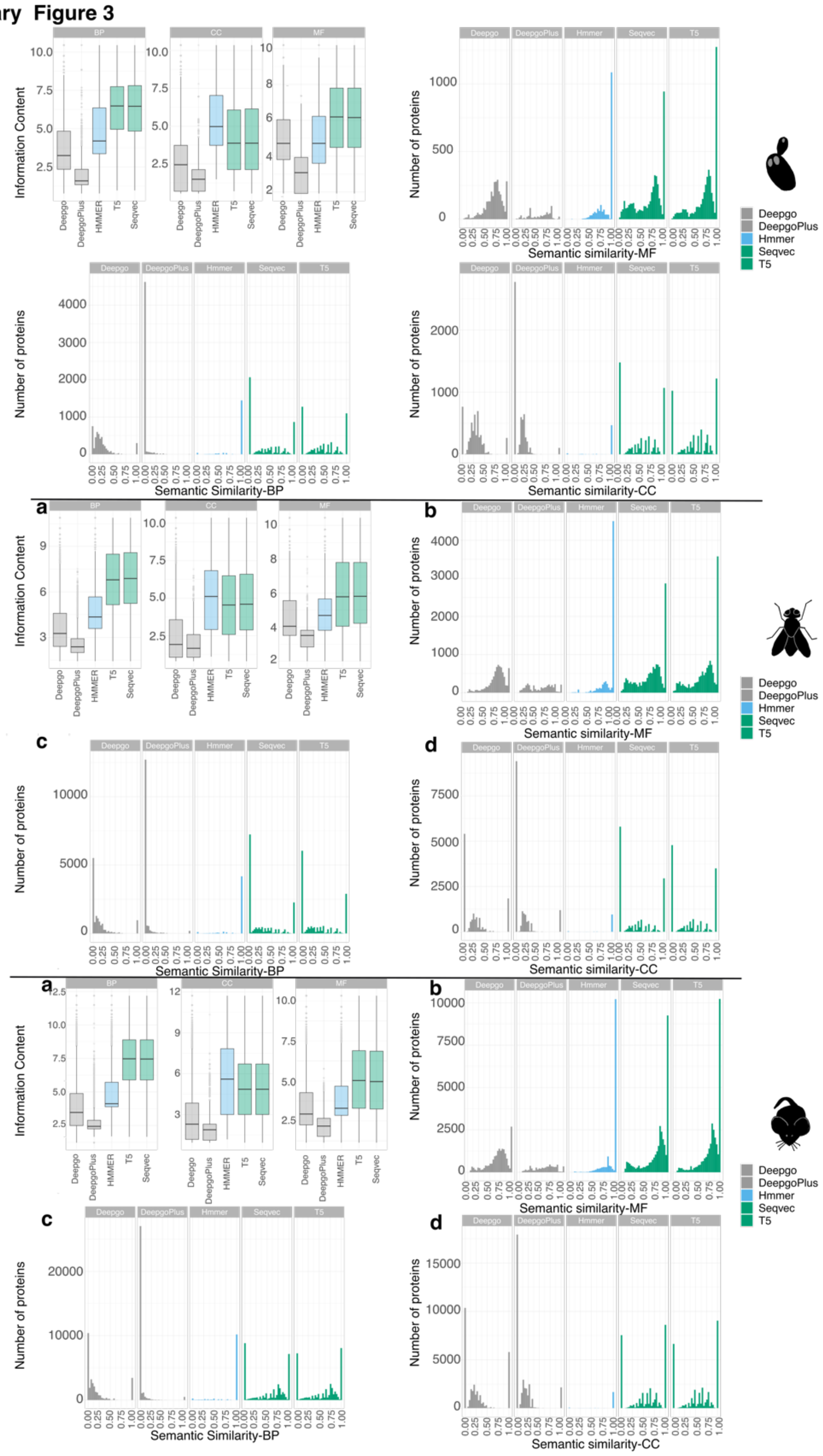


**Supplementary Figure SF3.** Information provided by the methods. **a.** Information content (IC, Y-axis) values for each method across ontologies: the larger the number, the more informative (more specific terms). **b-d**. The Semantic Similarity (SS) per protein (Y-axis) reflects the protein counts, the X-axis indicates the SS value with 1 indicating that the annotations and predictions are identical in **b:** MF, **c:** BP, and **d:** CC.

Supplementary Figure 4

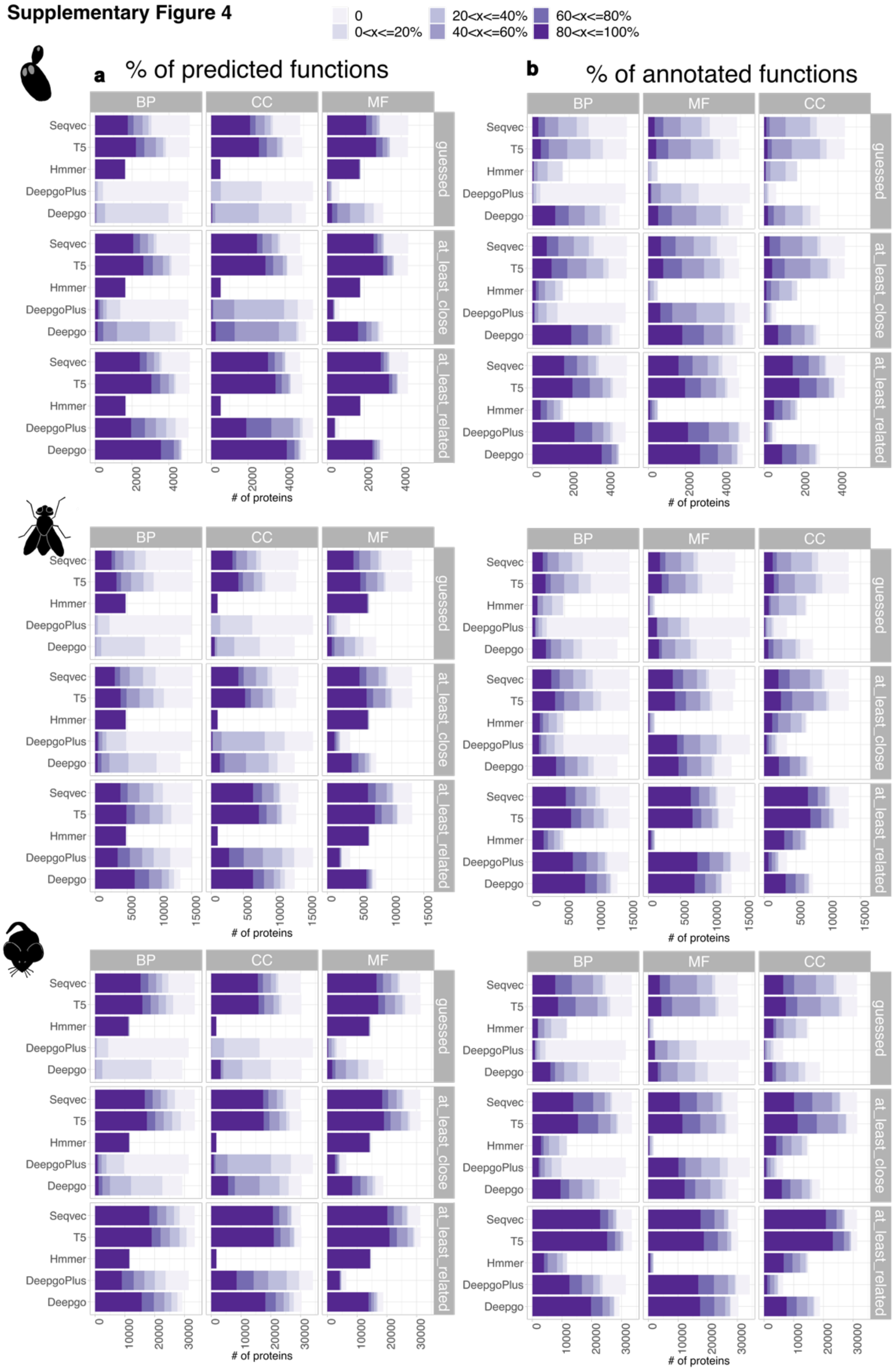


**Supplementary Figure SF4.** Per protein coverage of the methods. **a.** The proportion of correct predictions out of what could be predicted from the annotation space. **b.** represents how many are correct from the entire prediction space (are annotated). The colours indicate the percentage ranges.

Supplementary Figure 5

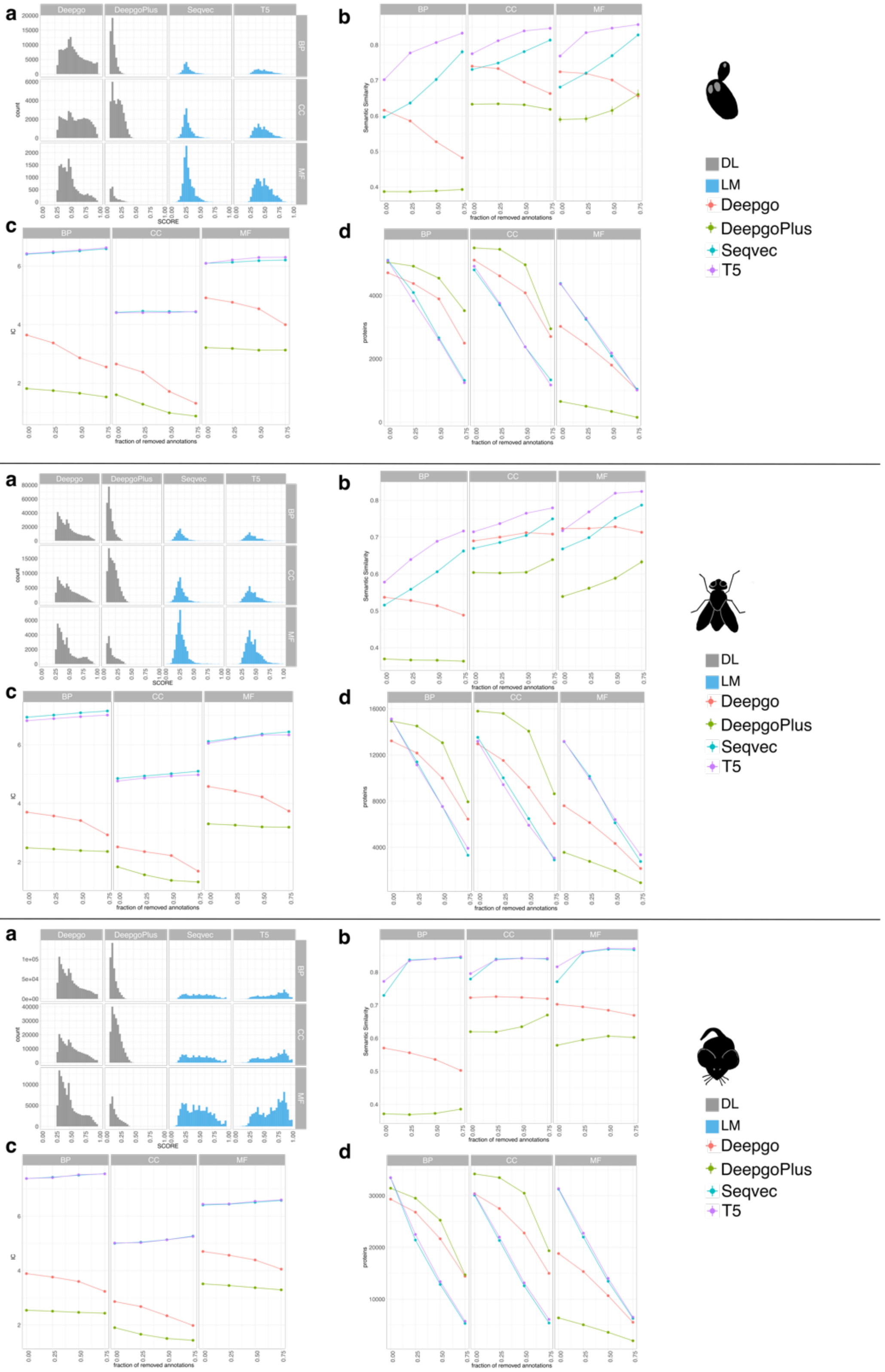


**Supplementary Figure SF5.** Score effect. **a.** Score distribution for each method in *C. elegans.* **b.** The effect of score on Semantic similarity (SS). **c.** The effect of score on information content (IC). **d.** The effect of score on protein coverage.

### Supplementary Figure 6

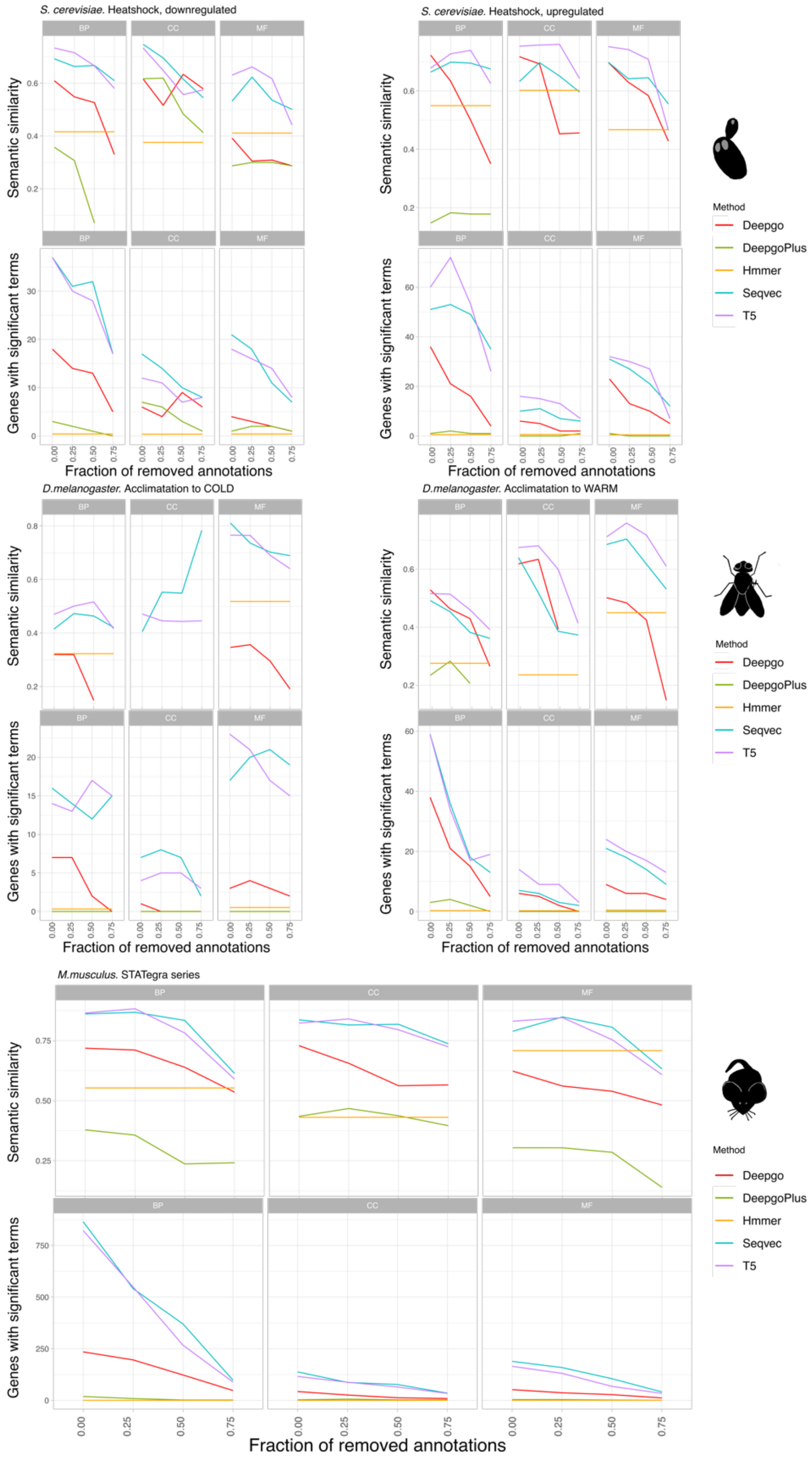


**Supplementary Figure SF6.** Recovery of functional information. Semantic similarity between GO terms from TopGO enrichment experiments, comparing the GO annotations in UniProt to the GO predictions: Upper panels, SS across score quantiles; Lower panels, number of proteins harbouring significant terms across quantiles.

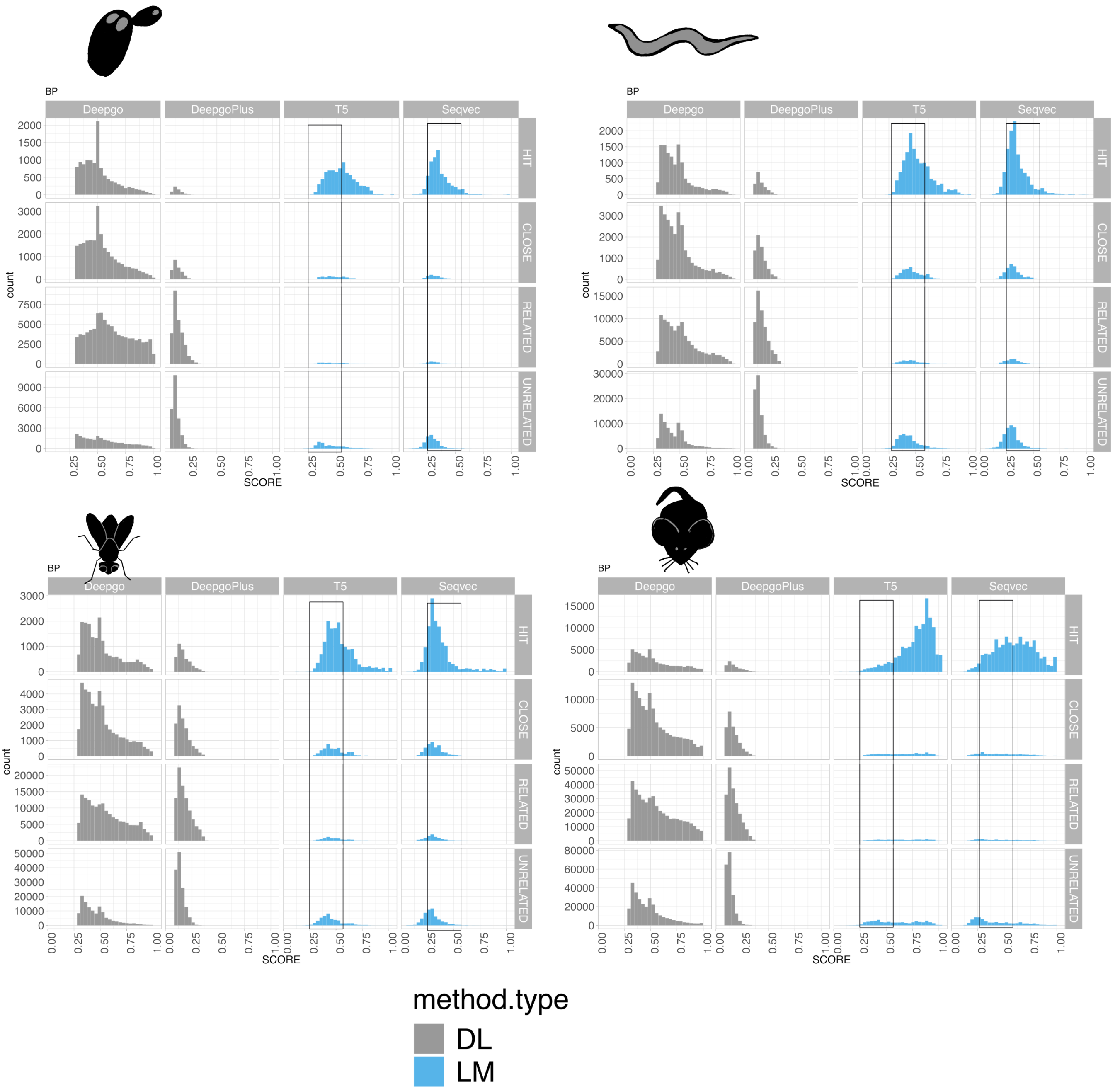


**Supplementary Figure SF7. Score distribution of our precision categories for the Biological Process ontology.** Numbers of Y axis are annotation counts, while numbers in the X axis indicate the fraction of removed annotations from the previous quantile. The boxed areas in the LM distributions indicate the counts from 0.25-0.50, since LMs are the only ones that cover the full range of possibilities (0-1). Overall, we observed a variation in the proportion of annotation counts at different scores for all organisms, except for in mouse where a displacement of the curve was observed to the right, indicating a potential influence from sister species

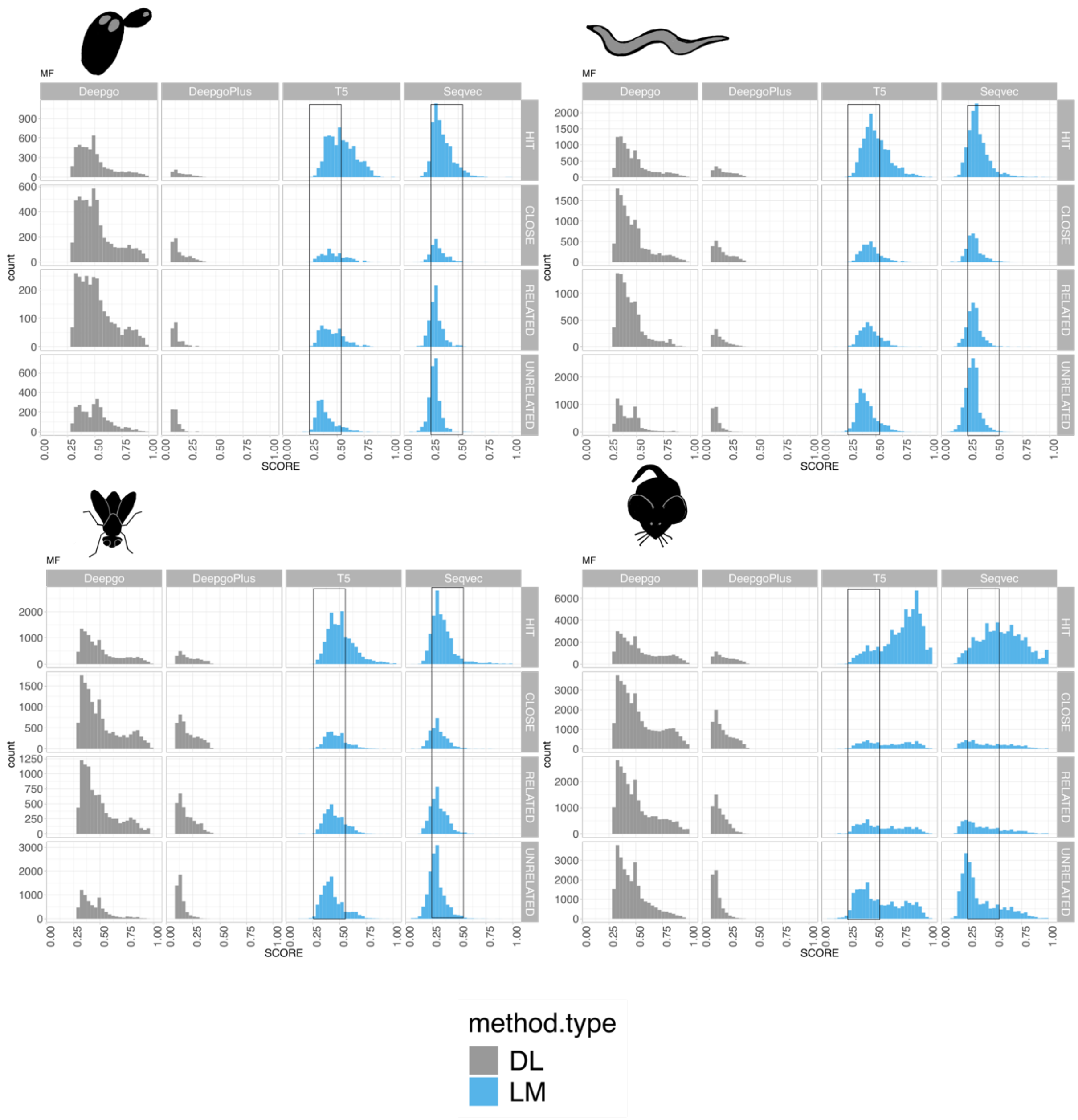


**Supplementary Figure SF8. Score distribution of our precision categories for the Molecular Function ontology.** Numbers of Y axis are annotation counts, while numbers in the X axis indicate the fraction of removed annotations from the previous quantile. The boxed areas in the LM distributions indicate the counts from 0.25-0.50, since LMs are the only ones that cover the full range of possibilities (0-1). Overall, we observed a variation in the proportion of annotation counts at different scores for all organisms, except for in mouse where a displacement of the curve was observed to the right, indicating a potential influence from sister species.

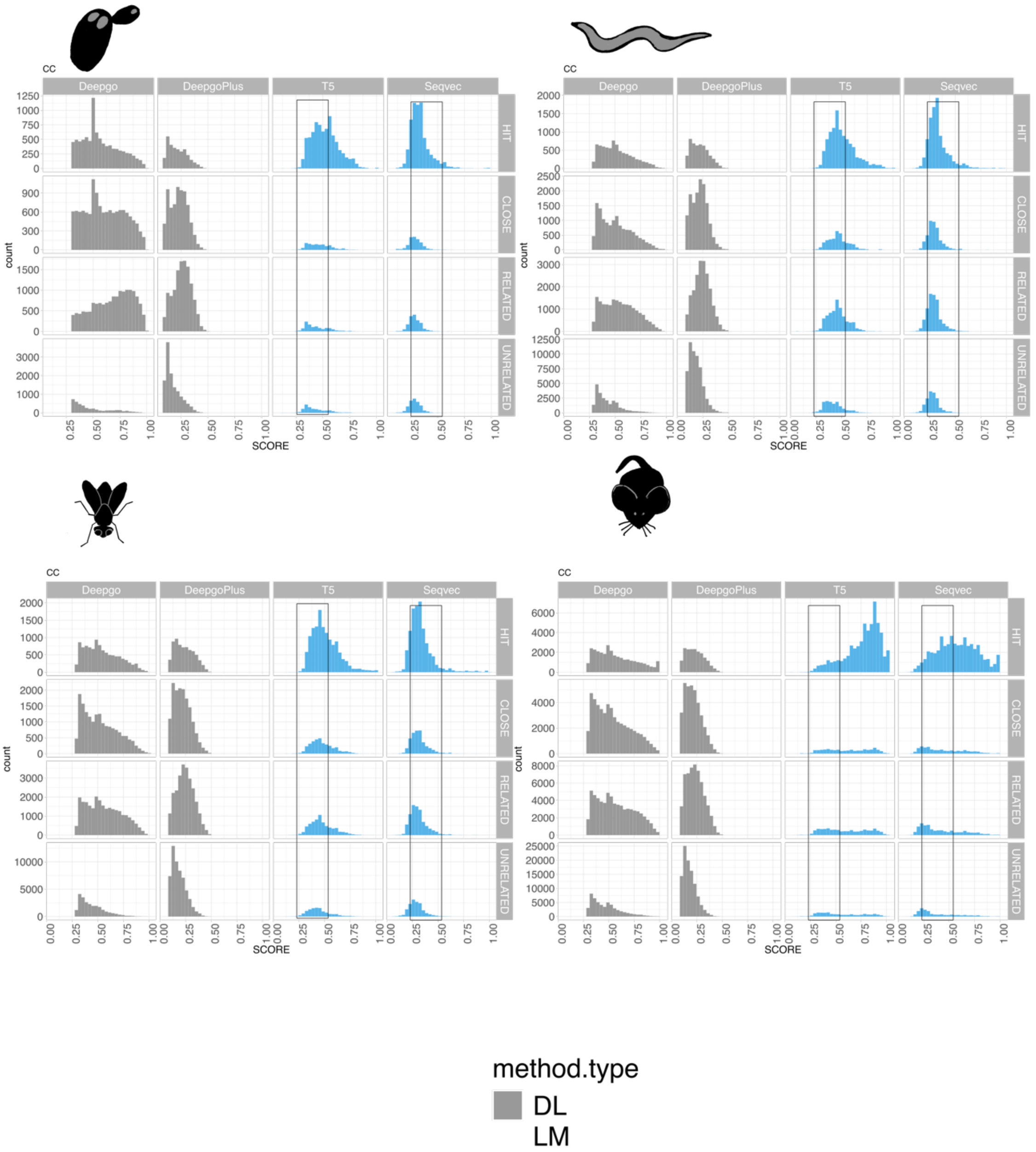


**Supplementary Figure SF9. Score distribution of our precision categories for the Cellular Component ontology.** Numbers of Y axis are annotation counts, while numbers in the X axis indicate the fraction of removed annotations from the previous quantile. The boxed areas in the LM distributions indicate the counts from 0.25-0.50, since LMs are the only ones that cover the full range of possibilities (0-1). Overall, we observed a variation in the proportion of annotation counts at different scores for all organisms, except for in mouse where a displacement of the curve was observed to the right, indicating a potential influence from sister species.
